# An *in vivo* knockdown strategy reveals multiple functions for circMbl

**DOI:** 10.1101/483271

**Authors:** Nagarjuna Reddy Pamudurti, Vinay Vikas Konakondla-Jacob, Aishwarya Krishnamoorthy, Reut Ashwal-Fluss, Osnat Bartok, Stas Wüst, Katerina Seitz, Roni Maya, Noam Lerner, Ines Lucia Patop, Silvio Rizzoli, Tsevi Beautus, Sebastian Kadener

## Abstract

Circular RNAs (circRNAs) are highly abundant and evolutionary conserved RNAs of mostly unknown functions. circRNAs are enriched in the brain and accumulate with age in flies, worms and mice. Despite their abundance, little is known about their functions, especially in the context of whole organisms. Here we report the development and use of shRNAs to knock down and study the function of circMbl, the most abundant circRNA in *Drosophila.* This circRNA is highly conserved through evolution and is generated from the locus of the essential splicing factor *muscleblind* (*mbl*). Briefly, we generated flies in which circMbl is reduced more than 90% without measurable off-target effects in the hosting gene as well as in other RNAs. These flies display specific defects that suggest roles of circMbl in muscle and neural tissues during development and in adult flies. More specifically, whole organism downregulation of circMbl leads to male developmental lethality, altered gene expression, behavioral defects, wing posture- and flight defects. Moreover, these phenotypes are recapitulated by a second shRNA targeting circMbl. Importantly, knockdown and overexpression of circMbl affect mostly the same genes but in the opposite direction. Last but not least, downregulation of circMbl in the fly central nervous system caused abnormal synaptic function. Together, our results demonstrate the functionality of circMbl at the organismal level likely by acting in multiple tissues. Moreover, here we provide the first proof of functionality of circRNAs in *Drosophila* as well as a methodological approach that enables the comprehensive study of circRNAs *in vivo*.

**SIGNIFICANCE STATEMENT:** Circular RNAs (circRNAs) are highly abundant and evolutionary conserved RNAs of mostly unknown functions. Here we report the development and use of a shRNA-based system to knockdown specific circRNAs *in vivo*. We generated flies in which circMbl, the most abundant circRNA is reduced more than 90% without measurable off-target effects. These flies display male developmental lethality, altered gene expression, behavioral defects, wing posture- and flight defects. These phenotypes are recapitulated by a second shRNA targeting circMbl. Moreover, downregulation of circMbl in the fly central nervous system caused abnormal synaptic function. Together, our results demonstrate the functionality of circMbl at the organismal level and provide a methodological approach that enables the comprehensive study of circRNAs *in vivo*.

## INTRODUCTION

Exonic circular RNAs (circRNAs) are a highly abundant type of RNAs produced through circularization of specific exons in a process known as back-splicing (1). circRNA biosynthesis is promoted by the presence of complementary sequences in flanking introns and/or by specific splicing factors like MUSCLEBLIND (MBL), QUAKING, and others (2-7). circRNAs are expressed in tissue- and development stage-specific ways, independently of the expression of the hosting gene (8). Indeed, abundant circRNAs can control gene expression in *cis* by competing for production with their hosting genes (6). Some circRNAs also produce proteins. The circRNA-encoded peptides usually share a start codon with the hosting genes and might be important in synapses and muscle (9-11).

Recent work has uncovered a handful of circRNAs that function in *trans*. For example, the circRNAs derived from CDR1as and *sry* likely regulate miRNA function and/or localization (12, 13). In addition, other circRNAs titrate or transport proteins and might be important for cancer development (6, 14, 15). circRNAs can also mediate responses to viral infections (16, 17). circRNAs are particularly enriched in neural tissue (18-21). Moreover, circRNA levels increase with age in the brains of mice and flies as well as in worms (21-23) and are affected by neuronal activity (18). These observations suggest important roles for circRNAs in the brain. Indeed, knockout of CDR1as in mouse results in abnormal gene expression in the brain and specific behavioral defects (24).

The *muscleblind* (*mbl*) locus of *Drosophila* encodes a highly conserved splicing factor important for muscle development and function (25). Indeed, the presence of CTG repeats in specific genes leads to the sequestration of the human homolog of MBL (MBNL1) which provokes myotonic dystrophy (26, 27). MBNL1 regulates splicing, 3′ end formation, and localization of many mRNAs, (28-30) and sequestration of MBNL1 results in dysregulation of expression of these mRNAs (31, 32). Importantly, MBNL1 has functions in other tissues. For example, MBNL1 is also expressed in the brain, where it has overlapping function with another protein from the MBNL family, MBNL2. MBNL1 and 2 regulate many aspects of RNA metabolism in the brain (28, 29, 33). Similar functions are carried out by the *Drosophila* MBL protein, which is expressed in the fly brain and muscle. *Drosophila* contains only one *muscleblind* homolog, MBL (34-36). Interestingly, the *muscleblind* gene also host the most abundant circRNA in *Drosophila* (circMbl) and recent work show that the production of this circRNA in flies and humans is driven by the MBL protein itself (6). Despite the abundance and evolutionary conservation little is known about the putative functions of circMbl. Tackling this issue is difficult due to the essential functions of MBL during development and in adult flies.

Here we report the development and use of short hairpin RNAs (shRNAs) to investigate putative functions of circMbl *in vivo.* Using this approach, we generated flies in which circMbl, is reduced more than 90% without measurable off-target effects. These flies display male developmental lethality, altered gene expression, behavioral defects, wing posture- and flight defects. These phenotypes are recapitulated by a second shRNA targeting circMbl. Moreover, downregulation of circMbl in the fly central nervous system caused abnormal synaptic function. Together, our results demonstrate the functionality of circMbl at the organismal level and provide a methodological approach that enables the comprehensive study of circRNAs *in vivo*.

## RESULTS

### circMbl can be specifically downregulated *in vivo* by miRNA-derived shRNAs

To determine functions of circMbl *in vivo* we expressed shRNAs directed against circMbl-specific back-spliced junction (Fig. 1A). We utilized a vector based on a miRNA-like precursor (miR-1 (37)). We first tested this approach in *Drosophila* S2 cells. We co-transfected cells with a plasmid containing a minigene promoting the expression of circMbl (6) and a control plasmid or a plasmid expressing the shRNA directed against the circMbl junction. Expression of the shRNA reduced the levels of circMbl by 5-fold (Fig. S1A).

**Figure 1.**
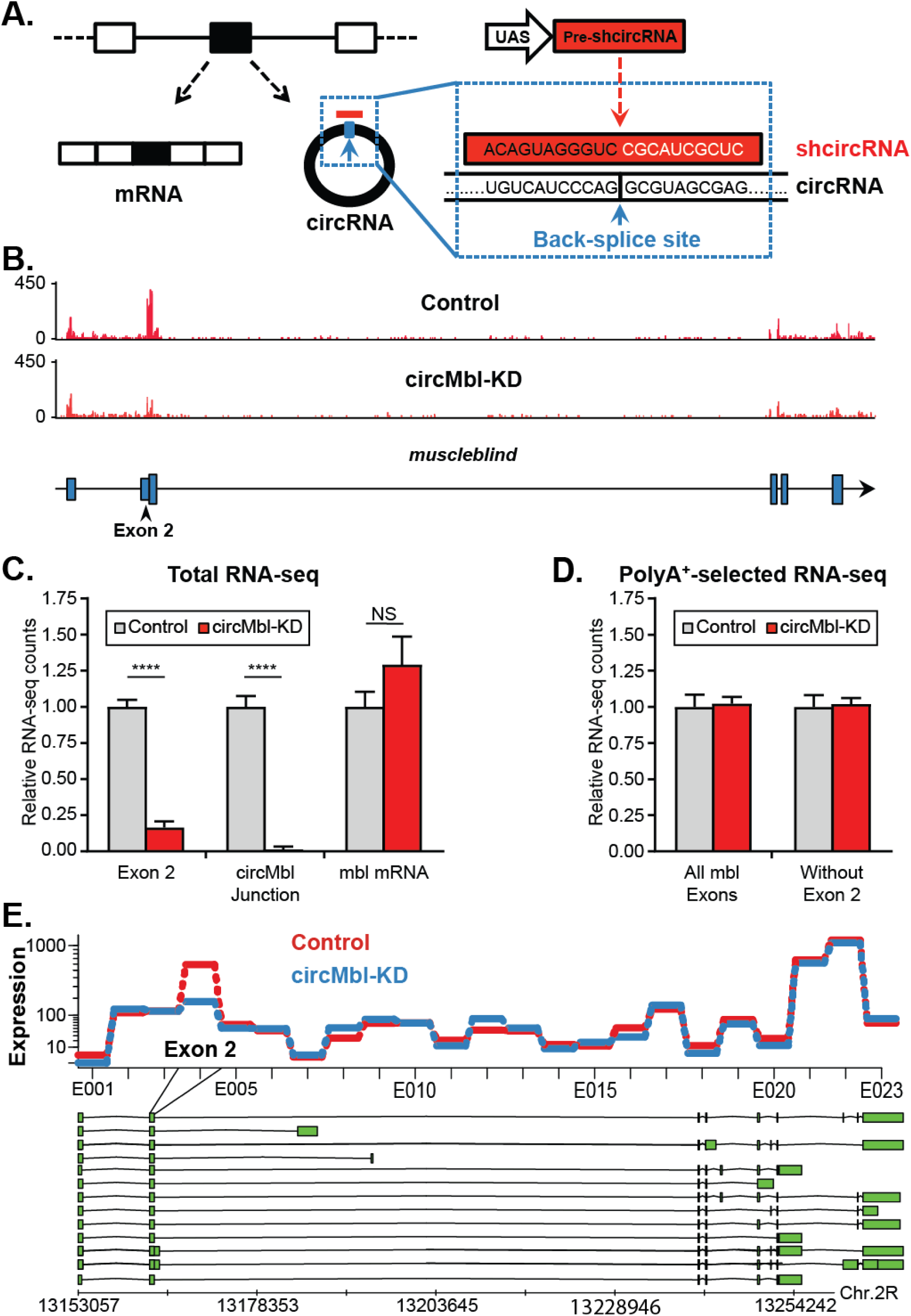
circRNAs can be downregulated by the use of miRNA-like shRNAs. **A.** Scheme of the knockdown strategy**. B.** IGV snapshot from total-RNA-seq data, showing a specific reduction of exon2 of *muscleblind* in respect to a control strain (UAS-shcircMbl), demonstrating a circRNA specific downregulation. The y-axis indicates the normalized RNAseq reads. **C.** Total-RNA-seq data quantification shows strong and specific circMbl reduction and no effect on the host *muscleblind* gene (the remaining exon2 reads derive from the linear gene (Student’s t-test; n=3; ****p<0.0001)). **D.** *Muscleblind* mRNA expression is preserved (PolyA^+^- RNA-seq; Student’s t-test; n=3). **E.** shRNA against *muscleblind* is not affecting alternative splicing of other *mbl* isoforms expression. Data presented is differential exon usage analysis (DEXseq, (44)).

We then utilized this plasmid to generated flies expressing the shRNA against circMbl. We analyzed gene expression by total (rRNA-) and polyA+ RNA-seq from controls and from flies expressing the shRNA under the control of a constitutive driver (*actin*-*Gal4*; circMbl-KD flies). We observed a specific and strong reduction in circMbl levels in fly heads (Fig. 1B and C). The effect was highly specific for the circular molecule as we did not observe a reduction of linear *mbl* mRNA (Fig. 1C and D). To rule out effects on specific *mbl* mRNA isoforms, we compared the levels of all known *mbl* isoforms and observed no significant differences between the circMbl-KD and control flies (Fig. 1E), demonstrating that the shRNA does not target any of the linear *mbl* mRNA isoforms.

### shRNA for circMbl does not display identifiable off-targets effects

Although the expression of the shRNA does not influence the levels of any of the known linear *mbl* transcripts, there is a possibility that it might alter their translation. Therefore, we compared the MBL protein expression in control flies to the circMbl-KD line, without detecting a difference (Fig. 2A).

**Figure 2.**
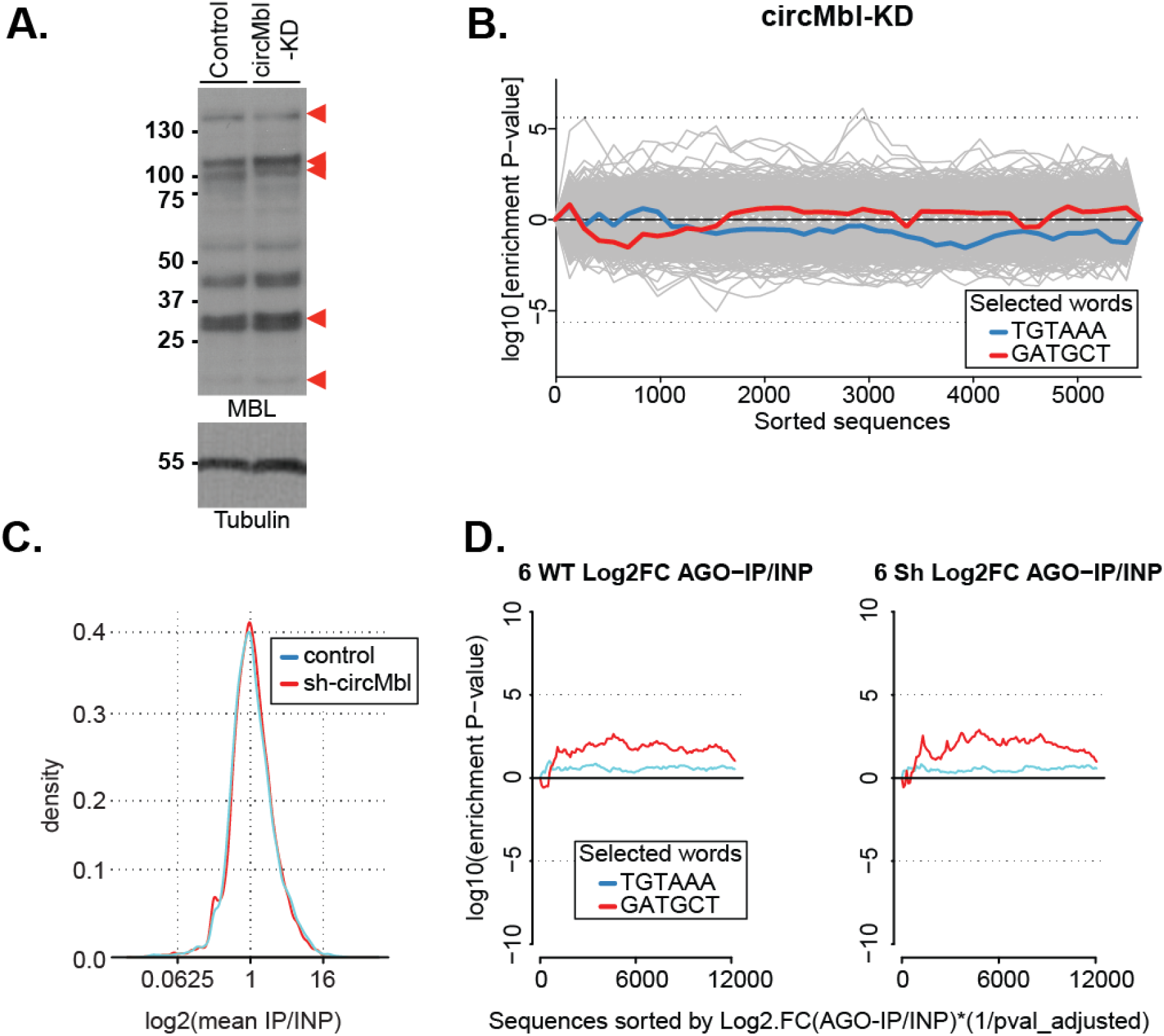
Knockdown of circRNAs is highly specific and does not affect the expression of the mRNA from the host gene. **A.** circRNAs KD does not affect the host gene protein expression. Western blot analysis of fly head extracts from circMBL KD, and controls. Red arrows represent known MBL isoforms in the anti-MBL blot. Tubulin is blotted as a loading control. Flies heads were utilized for preparing protein extracts. **B.** Assessment of off-targets by Sylamer. Traces show the seed enrichment for the genes differentially express upon downregulation of circMbl. shRNA and shRNA*-seed sequences shown in blue and red, respectively. For circMbl knockdown we did not observed any seed enriched among the downregulated genes. **C.** Expression of sh-circMbl does not alter the general binding of mRNAs to AGO-1. The panel shows the kernel density plot for the mean of the enrichment for each gene. **D.** Sylamer enrichment landscape plot for sh-circMbl and sh-circMbl* 6mers. The x-axis represents the genes sorted from the most to the least enriched in the AGO-1 IP-seq. The y-axis shows the hypergeometric significance for each word at each leading bin.

However, it is possible that shRNAs could target mRNAs with more restricted base complementarity. To test this possibility, we identified mRNAs with more than twelve bases complementarity to the shRNA against circMbl using nucleotide blast for Drosophila melanogaster (taxid:7227). We found 6 transcripts (beside mbl) that potentially could be affected (Table S1). After analysis of the transcriptome data we found none of the transcripts significantly reduced (Table S2).

The shRNA could also have off-target effects on other mRNAs by acting as miRNAs. Therefore, we determined whether the downregulated mRNAs in shRNA-expressing strain are enriched for the seed of the shRNA or shRNA* utilizing the SYLAMER algorithm (38). None of the downregulated mRNAs enriched for any of the relevant seed sequences (Fig. 2B).

Although unlikely, it is still possible that the shRNAs could perturb generally the miRNA-regulated mRNAs or provoke changes in translation of mRNAs without significant effects at the RNA and protein levels. To rule out this possibility, we then determined how the expression of the shRNA against circMbl alters the population of mRNAs bound to AGO1, the key component of the miRNA effector machinery in *Drosophila* {Forstemann, 2007 #838}. To do so we sequenced mRNAs which co-purify with AGO1 in heads of control and circMbl-KD flies. We utilized a similar approach as previously described (39). As expected mRNAs bound to AGO1 are highly enriched for mRNAs containing predicted miRNA binding sites (Fig. S1B). Importantly, expression of the shRNA against circMbl did not alter the general profile of binding of RNAs to AGO1 (Fig. 2C), demonstrating that expression of the shRNA does not flood AGO1 containing complexes and strongly suggesting that it is not even loaded to AGO1. Moreover, the kmer enrichment found on the AGO-1 bound mRNAs for both control flies and circMbl KD flies is similar and it does not show significant enrichment for the 6mers that could be generated from the-processing of sh-circMbl oligonucleotide (Fig. 2D). Furthermore, the few mRNAs which were differentially bound to AGO1 in the circMbl KD flies were not enriched for seed sequences potentially targeted by the shRNA and shRNA* in this strain (Fig. S1C), demonstrating that the shRNAs does not bind and modulate other RNAs. All the results presented above demonstrated that the shRNA for circMbl is highly specific and suitable for determining the functionality of this circRNA *in vivo*.

### Modulation of circMbl alters expression of brain and muscle-related genes

We recently generated flies expressing a UAS-circMbl minigene, which allows overexpression (OE) of circMbl (9). To identify genes affected by modulation of circMbl we generated and sequenced 3′-RNA-seq libraries from heads of control, circMbl-KD, and circMbl-OE flies. We then identified 39 mRNAs that are differentially expressed both in circMbl-KD and circMbl-OE flies when compared to the corresponding control strain (Fig. 3A, Table S3). Four of these mRNAs had similar trends in both strains, suggesting that these genes might be related to general transgene expression. 35 genes however, showed opposite trends in the OE and KD strains (Fig. 3A). Strikingly, this group of genes is enriched for genes involved in muscle development and function (Fig. S2). In addition, we found that muscle- and flight-related GO terms were significantly enriched for genes differentially expressed in the circMbl OE strain (Fig. S2), suggesting that circMbl function is related to muscle and flight.

**Figure 3.**
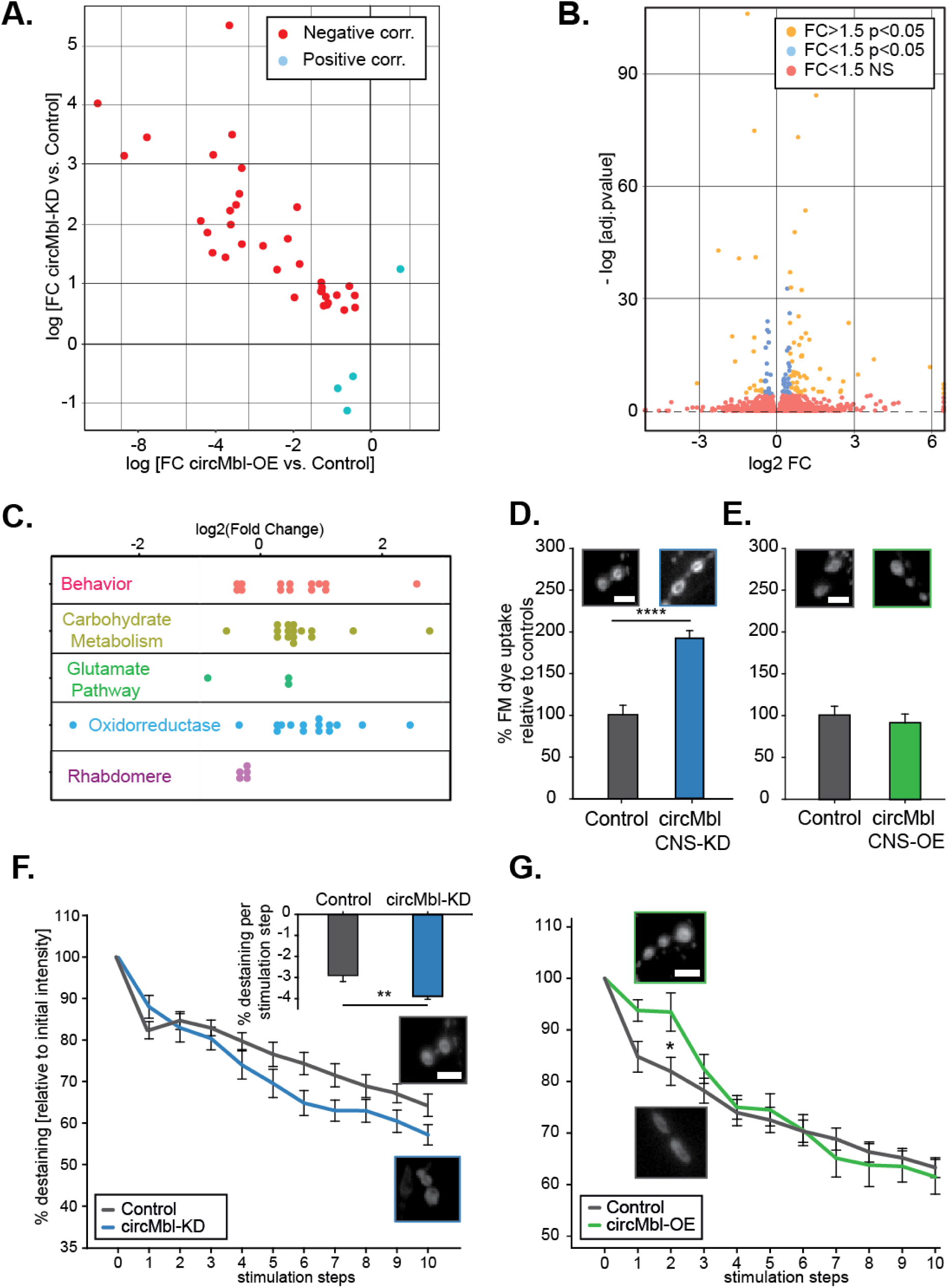
Knockdown of circMbl affects gene expression and synaptic function. **A.** Fold changes (in log scales) between circMbl OE (x-axis) and circMbl-KD (y-axis) in comparison with their respective controls. Blue indicates genes showing positive correlation in both comparisons. In red, the genes showing negative correlation between the fold change in one comparison and the other. In all cases we considered genes with fold change > 1.5 and p-value<0.05. **B.** Volcano plot showing gene expression differences in fly heads between control and *elav*-Gal4:UAS-Dcr2/UAS-shcircMbl (CNS-KD) flies. Orange dots indicate RNAs with fold change >1.5 and p-value<0.05. **C.** Gene Ontology (GO) terms significantly enriched among genes differentially expressed between the heads of control and circMbl CNS-KD flies. Dots indicate individual genes. Adjusted p-value<0.05. **D.** FM dye fluorescence in boutons of 3^rd^ instar larval motoneurons, following high potassium stimulation in presence of 10 μM FM 1-43. Fluorescence intensities of boutons of circMbl CNS-KD animals (*elav*-Gal4, dcr2 x UAS-shMbl; cyan bar) are normalized to the mean fluorescence of boutons of control animals (elav-gal4, dcr2 x uas- shLuciferase; dark grey bar). The bars show mean and SEM (n≥14 boutons from at least four different animals; ****p<0.0001). Insets show exemplary images. Scale bar 2 μm. **E.** Similar than D. but comparing the boutons of control (*elav*-gal4; grey bar) with cirMbl CNS OE (*elav*-gal4; UAS-circMbl) flies. (n≥12 boutons from at least four different animals). **F.** Subsequent to FM dye loading, preparations were electrically stimulated in a step-wise fashion (10 trains at 20 Hz for 10 seconds each). The line graph shows the fluorescence loss relative to the initial fluorescence intensity for each bouton (grey dots indicate controls, cyan dots indicate circMbl knockdown). Images show representative images. The inset indicates the relative destaining amount of the individual boutons per stimulation step (−2.9% for control, −3.9% for circMbl knockdown), as determined by linear fits to the individual fluorescence curves (**p=0.006). Data are mean and SEM of n≥14 boutons from at least four different animals. **G.** Similar than F but for circMbl CNS OE and control strains (defined as in F). *p=0.01 as determined by unpaired Student’s t-test for the difference in fluorescence loss between the conditions after the second stimulation step. Data are mean and SEM of n≥12 boutons from at least four different animals.

Our recent report suggested that circMbl might regulate MBL function (6). Interestingly, the function of MBNL1, a human orthologue of MBL, is strongly compromised in individuals with myotonic dystrophy (DM (27)). We therefore determined whether the genes modulated by circMbl are altered upon knockdown of *mbl* or in a fly model of DM (using an already published dataset (40)). Strikingly, we found that eight of the 39 genes affected by the circMbl manipulations were also altered in the *mbl* knockdown and DM flies (Table. S3). Moreover, six of these eight mRNAs are downregulated upon *mbl* knockdown and in the DM model and were upregulated in the circMbl-KD strain. These same six mRNAs were also downregulated upon circMbl overexpression and include genes related to flight and muscle function. These results support the notion that circMbl regulates MBL function, although we cannot rule out a more complexed regulation, as MBL also regulates circMbl production (6).

### Modulation of circMbl levels alters synaptic vesicular function

To knockdown circMbl in a more specific cellular population, we utilized the CNS-specific driver *elav*-Gal4. Using this driver, circMbl levels were reduced more than 80% in *Drosophila* heads (Fig. S3A). Co-expression of Dicer-2 enhanced the silencing of circMbl (Fig. S3A). In all cases the levels of linear *mbl* were not affected (Fig. S3B). A small number of genes were misregulated in circMbl-CNS-KD flies (Fig. 3B). Among these were genes involved in glutamate metabolism, behavior, and other brain-related processes including light processing and behavior (Fig. 3C).

To assess the importance of circMbl on the synaptic physiology of the *Drosophila* larval neuromuscular junction (NMJ), we monitored dye uptake and destaining in flies with downregulated or upregulated circMbl in the CNS. Briefly, we treated body wall muscle preparations of third instar *Drosophila melanogaster* larva with the dye FM 1-43 to load recycling vesicles of the NMJ synaptic boutons. We then washed and stimulated 10 times. We took images of the same synaptic boutons before stimulation and after each stimulation step. Quantification of the baseline fluorescence intensity of the boutons provides information about the size of the recycling pool of vesicles, whereas analysis of the fluorescence change over the course of the 10 stimulation steps allows conclusions to be drawn regarding the dynamics of exocytosis in response to stimulation.

Using this assay, we found that knockdown of circMbl resulted in significantly higher dye uptake relative to control (*p* < 1×10^-6^), indicating a larger pool of recycling vesicles (Fig. 3D). NMJ boutons of larva overexpressing circMbl showed dye uptake comparable to the control (Fig. 3E). We then analysed the destaining kinetics during stimulation (Fig. 3F and G). Boutons from circMbl overexpressing larva destained more slowly during the first few stimulation steps compared to controls but reached levels and rates comparable to controls by the 7^th^ to 8^th^ stimulation step. Boutons from circMbl knockdown flies displayed a stronger relative destaining per stimulation step than controls (*p* <0.006). This opposing effect suggest a role of circMbl in the regulation of the size of vesicle pools and release propensity of synaptic vesicles.

### Reduction of circMbl levels leads to developmental and adult phenotypes

circMbl-KD flies displayed male developmental lethality with high penetrance (see below). Moreover, a large proportion of the circMbl-KD males that escaped the developmental lethality displayed a strong wing-posture phenotype (Fig. 4A). Females of the same strain displayed a similar phenotype when raised at 29 °C (Fig. 4A). This temperature increases the efficiency of the Gal4 system and/or adds stress to the system during development. Wing posture phenotypes are generally related to defects in the indirect flight muscles or in the motor neurons that control these muscles. Similar wing phenotypes were described in a fly model of DM (41).

**Figure 4.**
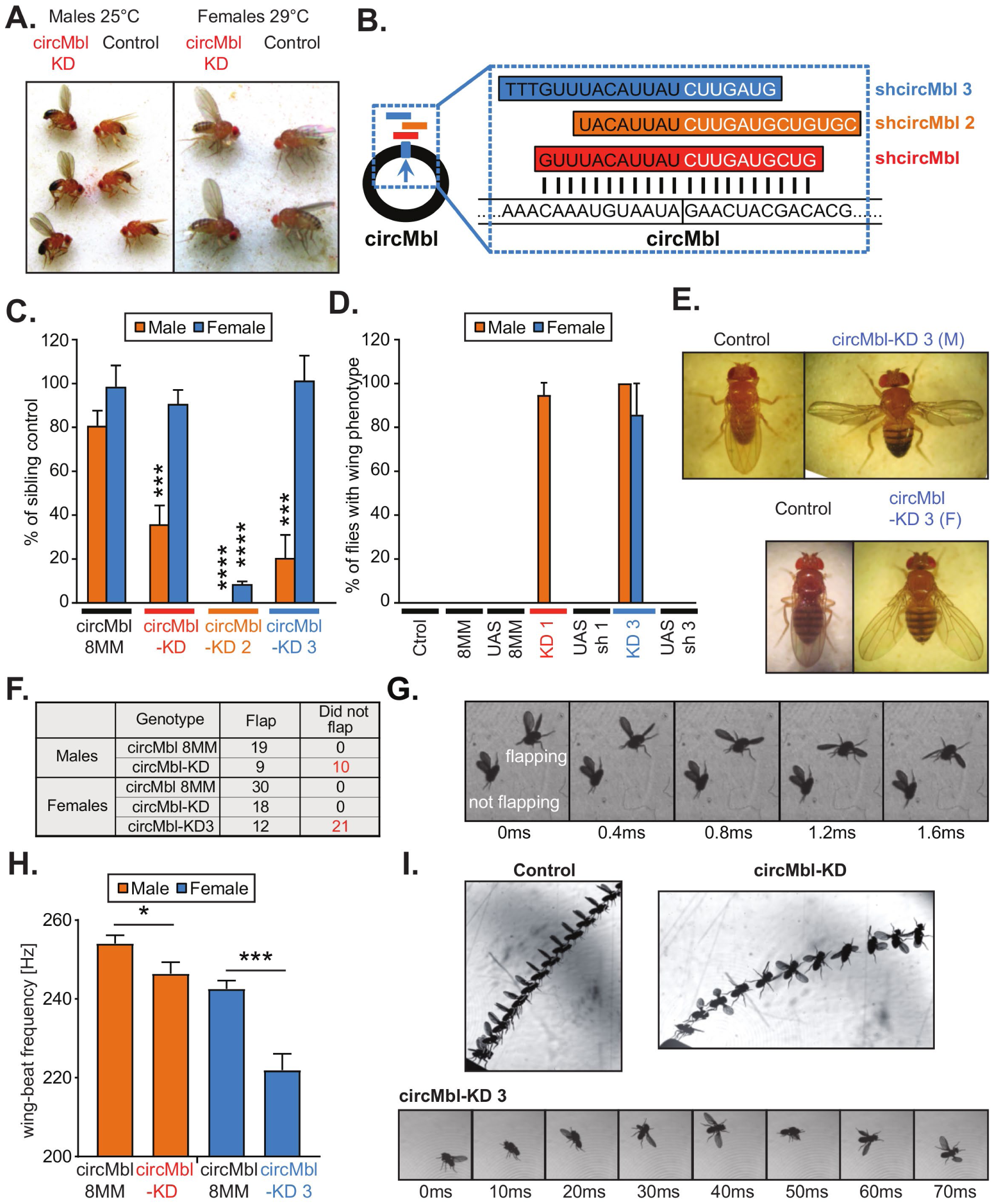
Knockdown of circMbl provokes male lethality and a characteristic wing posture. **A.** Picture of circMbl-KD males with “wings up phenotype” reared at 25 °C (left) or females (right) reared at 29 °C next to control flies (*actin*-Gal4) reared in the same conditions. **B.** Scheme of the additional shRNAs designed against circMbl. **C.** Male lethality of the control (*actin-*gal4, UAS- shcircMbl 8MM) and the indicated circMbl shRNAs. We plotted % of male and females against the sibling controls. N= 8 for circMbl-KD flies and 7 for the rest of the strains (comparison of KD lines to controls of same sex; Student’s t-test; ***p<0.0005, ****p<0.0001). **D.** Percentage of male and females presenting wing up (circMbl-KD) or open (circMbl-KD 3) phenotypes. 8 independent experiments for circMbl-KD flies and 7 for the other strains. **E.** Pictures of circMbl- KD 3 males (M) and females (F) next to control (*actin*-Gal4). **F.** Results of the tapping assay. **G**. A sequence of side-view images from the tapping assay taken 0.4 ms apart. All videos were taken in 10,000 frames/sec. Images show two male circMbl-KD flies falling side by side: the fly on the right flapped its wings while the fly on the left did not. **H**. Mean wing-beat frequency in the free- flight assay. We measured ~30 flies from first three lines and 12 flies from the circMbl-KD 3 (n=25/32/29/12; Student’s t-test; *p<0.05, ***p<0.0005). **I.** Representative flight events from the free-flight assay. Top-left: a male circMbl 8MM (Control) fly taking off normally. Superposed images are shown every 4 ms. Top-right: a male circMbl-KD fly taking off. Superposed images are shown every 6 ms. Bottom: a female circMbl-KD3 fly shown shortly after take-off. Images are shown every 10 ms. Although the fly seemed to take off normally, at 30 ms it went unstable and crashed, falling backwards.

To rule out non-specific effects of the shRNA, we generated two additional shRNAs against the circMbl junction. The new shRNAs are perfectly complementary to the circRNA, but target sites were shifted by three nucleotides in either direction with respect to that of the original shRNA (Fig. 4B).

Expression of any of the two shRNAs provoked an almost complete silencing of circMbl with no significant effect on the linear *mbl* transcript when compared to the 8MM control. (Fig. S4A). Because of the male lethality we measured the knockdown efficiency only in females. As control we generated a fly line expressing a shRNA with 8 mismatches to the circMbl junction (circMbl- 8MM).

When expressing the shRNAs under the constitutive control of the *actin-*Gal4 driver, we observed significant but incomplete developmental male lethality with one of the shRNAs (circMbl-KD3; Fig. 4C), similar to lethality levels observed with the originally tested shRNA (Fig. 4C). Expression of the other shRNA (circMbl-KD2) resulted in complete male lethality, and a very strong lethality in females (Fig. 4C). Importantly, expression of the shRNA with mismatches to the circMbl junction (circMbl-8MM) did not affect viability or the number of males (Fig. 4C). We observed normal wing postures in all control strains. However, similar to the originally tested circMbl-KD line, all circMbl-KD3 males and females displayed wing posture phenotypes (Fig. 4D and E). These results demonstrate that downregulation of circMbl using different shRNAs provoke related phenotypes.

We also tested whether downregulation of circMbl alters general motor function. To do so we determined the climbing ability of circMbl KD flies. Indeed, we found that circMbl-KD and circMbl-KD3 flies show severe defects in their climbing ability (Fig. S4 B - D) strongly suggesting roles of this circRNA in locomotion and muscle function. We did not perform this or any of the flight assays described below in circMbl-KD3 males as the few that eclosed died within the first few days.

The observed wing phenotypes and changes in gene expression suggest that circMbl is necessary for correct muscle function and flight. To test this hypothesis, we evaluated the flight of the different circMbl-KD strains using high-speed recordings. We first determined whether control, circMbl-KD, and circMbl-KD3 flies could flap their wings when released after tapping the bottle. In these conditions all the males and females from a control strain flapped their wings (Fig. 4F, Video S1). Similarly, all the female flies from the circMbl-KD strain flapped their wings (Fig. 4F, Video S2). Only half (9/19) of the males from the circMbl-KD strain and one-third (12/33) of circMbl-KD3 females managed to flap their wings (Videos S3 and S4 respectively, Fig. 4F for the summarized results, Fig. 4G for an example of males from circMbl-KD). These results show a direct correlation between the wing posture and the capacity of the circMbl flies to flap (as the circMbl-KD females grew in these conditions did not display wing phenotypes). Since the tapping assay introduces additional mechanical stress compared to free-flight assays, we performed a second type of assay in which we carefully placed the flies and allowed them to take off and fly freely. Using this free-flight assay, we determined the mean wing beat frequency per fly. Males of the circMbl-KD strain and females of the circMbl-KD3 strain displayed significantly lower wing- beat frequencies than controls (Fig. 4H and S4E). In addition, we observed that many of circMbl- KD3 females could not sustain normal flight and tended to lose flight stability and “crash” even after a seemingly normal take-off and while beating their wings (Fig. 4I). In sum, these physiological studies demonstrate a role for circMbl in flight, likely by modulating muscle function.

## DISCUSSION

Here we report the development and use of short hairpin RNAs (shRNAs) to investigate the functionality of circMbl *in vivo.* Using this approach, we generated flies in which circMbl is reduced more than 90% without measurable off-target effects. These flies display male developmental lethality, altered gene expression, behavioral defects, wing posture- and flight defects. Moreover, downregulation of circMbl in the fly central nervous system caused abnormal synaptic function.

Even though hypothetically the shRNA could provoke small changes in dozens of genes (by acting as a miRNA), our AGO1-seq experiment strongly suggest against this possibility. Although further studies (e.g. CRISPR-based loss of function alleles and/or rescue experiments) might be nice to further rule out potential side effects, we clearly demonstrate that there is no measurable evidence of off-targets effects of our system so far. Importantly, this system offers the possibility of performing knockdowns restricted in space and time in *Drosophila* using the GAL4-GAL80 system {McGuire, 2004 #839}. This would allow to determine the specific stages and cell types at which circRNA expression is essential for development.

After we ensured the functionality of our system, we analyzed the physiologic changes after the loss of circMbl. The most sever phenotype was the partial or complete male lethality that was observed in all shRNA-lines. The line with the complete male lethality also showed partial female lethality. These stronger effects are likely due to a stronger silencing of the RNA or maybe the linear *mbl* during development. Indeed, several phenotypes seems to be more severe in males. This could be due to technical issues (i.e. weaker Gal4 expression in females) or to specific regulation of male-relevant genes by circMbl. The latter seems more likely due to the extensive sex-specific splicing regulation described in *Drosophila* The surviving individuals had severe defects for performing different motor tasks like climbing and flying. These defects are likely due to overlapping functions of the circRNA in muscle and brain tissues.

While the experiments presented above demonstrate the functionality of circMbl, they provide little insight into the molecular mechanism by which this circRNA operates. We previously demonstrated that MBL regulates circMbl production (6). Moreover, we found that MBL strongly binds to circMbl and we proposed that circMbl might regulate MBL production, function and/or localization. However, we did not see changes in *mbl* RNA and protein upon circMbl knockdown, suggesting that the interplay between circMbl and MBL happens mainly at the biogenesis level. It is also possible that circMbl regulates MBL localization or function but further experiments are needed to test this possibility. In this context, the circMbl-KD flies constitute an excellent system to determine how circMbl potentially alters MBL function *in vivo.* Moreover, given the role of MBNL1 in myotonic dystrophy, it will be interesting to determine whether circMBNL1 has any contribution to the development of the disease. The changes we observed upon modulation of circMbl in flies suggest this might be the case. Briefly we identified the same set of muscle related genes regulated in the circMbl knockdown flies and the Drosophila DM model. While the set of six muscle genes was reduced in the DM model, the knockdown of circMbl led to an increased expression of these genes. The overexpression of circMbl caused a decrease. These data suggest a complex but important relationship between MBL and circMbl for muscle function and a potential impact on the development of DM in the fly model. Interestingly, the phenotypes observed upon circMbl knockdown do not directly correlate with those previously described while modulating MBL protein. The latter include total developmental lethality, wing (42) and eye (43) development phenotypes. All the above suggest that while circMbl and MBL functions are tightly interconnected, the mechanism of interaction between these molecules is complex.

## Supporting information

## Acknowledgments

This work was funded by the European Research Council Consolidator Grant (ERC#647989) to SK, the Israel-Niedersachsen grant to SK and SR and NIH R01 grant (R01GM122406) to SK. All the RNA-seq data has been submitted to GEO (entries GSE122693, GSE122694 and GSE118360).

## Author Contributions

NRP generated the collection and performed most of the physiological assessments; VVK and AK performed the behavioral assays; RAF and ILP performed the computational analysis; OB generated the libraries and performed the western blots; SW helped with the figures and manuscript, KS and SR performed the NMJ synaptic experiments; RM, NL and TB performed the flying assays, and SK designed the experiments and wrote the manuscript.

## METHODS

### Fly strains and reagents

#### Fly strains

Wild type flies that we used in this study are *yw* and w1118 strains (Bloomington *Drosophila* Stock Center Indiana, USA). *Elav*-Gal4; UAS *Dcr2* were generated by using *elav*-Gal4 (stock number 458, Bloomington *Drosophila* Stock Center, Indiana, USA) and UAS-*Dcr2* flies. The circMbl OE strain is described in (9). Unless indicated otherwise, all crosses were performed and raised at 25 °C.

#### Generation of shRNA lines

To generate circMbl KD flies we designed oligonucleotides with perfect 21-nucleotide complementary sequence to the circRNA junction (or modification as stated in the text), annealed them, and ligated in to the linearized Valium20 vector with EcoR1 and Nhe1 restriction enzymes. Colonies were screened by PCR and the plasmid was purified and sequenced from positive colonies. These plasmids were sent for injection to BestGene Inc (CA, USA). Oligonucleotides used are annotated in Table S4. Potential off-targets were verified by Blast against the fly genome and transcriptome.

### Molecular Biology Methods

#### Cell culture and transfections

*Drosophila* S2 cells were cultured in 10% fetal bovine serum (Invitrogen) insect tissue culture medium (Biological industries). Transfection was performed at 60-80% confluence according to manufacturer protocol with TransIT2020 (Mirus Bio, MIR5400A) with a total of 2ug DNA in a six-well plate. In shMBL experiments 500ng circMBL plasmid, 250ng UAS-shMBL plasmid and 25ng of pActin Gal4 plasmid were used. For copper (Cu) induction 500μM copper were added to the media and after 24h cells were collected.

#### Real Time PCR analysis

Total RNA was extracted from adult fly heads using TRI Reagent (Sigma) and treated with DNase I (NEB) following the manufacturer’s protocol. cDNA was synthesized (using iScript and random primers, Bio-Rad) and used for qRT-PCR with the C1000 Thermal Cycler Bio-Rad and SYBR green, Bio-Rad. Cycling parameters were 95 °C, 3 min, 40 cycles of 95 °C, 10 s, 55 °C, 10 s and 72 °C, 30 s. The sequences of all the primers used in this assay are detailed in Table S4.

#### RNA libraries preparation for RNA-seq analysis

All information about the RNA-sequencing samples and types of libraries is detailed in Table S5. Total RNA was extracted using Trizol (Sigma) and treated with DNase I (NEB) following the manufacturer’s protocol.

Stranded ligation-based, **total-RNA libraries** preparation was modified from (Engreitz et al., 2013) as follows**: For PolyA+ libraries,** 0.5μg of total RNA was polyA+ selected (using Oligo(dT) beads, Invitrogen), fragmented in FastAP buffer (Thermo Scientific) for 3min at 94°C, then dephosphorylated with FastAP, cleaned (using SPRI beads, Agencourt) and ligated to a linker1(5Phos/AXXXXXXXXAGATCGGAAGAGCGTCGTGTAG/3ddC/, where XXXXXXXX is an internal barcode specific for each sample), using T4 RNA ligase I (NEB). Ligated RNA was cleaned-up with Silane beads (Dynabeads MyOne, Life Technologies) and pooled into a single tube. RT was then performed for the pooled sample, with a specific primer (5′-CCTACACGACGCTCTTCC-3′) using AffinityScript Multiple Temperature cDNA Synthesis Kit (Agilent Technologies). Then, RNA-DNA hybrids were degraded by incubating the RT mixture with 10% 1M NaOH (e.g. 2ul to 20ul of RT mixture) at 70°C for 12 minutes. pH was then normalized by addition of corresponding amount of 0.5M AcOH (e.g. 4ul for 22 ul of NaOH+RT mixture). The reaction mixture was cleaned up using Silane beads and second ligation was performed, where 3′end of cDNA was ligated to linker2 (5Phos/AGATCGGAAGAGCACACGTCTG/3ddC/) using T4 RNA ligase I. The sequences of linker1 and linker2 are partially complementary to the standard Illumina read1 and read2/barcode adapters, respectively. Reaction Mixture was cleaned up (Silane beads) and PCR enrichment was set up using enrichment primers 1 and 2: (5’AATGATACGGCGACCACCGAGATCTACACTCTTTCCCTACACGACGCTCTTCCGA TCT-3′, 5’-CAAGCAGAAGACGGCATACGAGATXXXXXXXXGTGACTGGAGTTCAGAC GTGTGCTCTTCCGATCT-3′, where XXXXXXX is barcode sequence)

and Phusion HF MasterMix (NEB). 12 cycles of enrichment were performed. Libraries were cleaned with 0.7X volume of SPRI beads. Libraries were characterization by Tapestation. RNA was sequenced as paired-end samples, in a NextSeq 500 sequencer (Illumina).

**rRNA- libraries** were similarly prepared, without the polyA+ selection step: 0.25μg of total RNA from each sample were fragmented in FastAP buffer (Thermo Scientific) for 3min at 94°C, then dephosphorylated with FastAP, cleaned and ligated to linker1 using T4 RNA ligase I (NEB). Ligated RNA was cleaned-up with Silane beads (Dynabeads MyOne, Life Technologies) and pooled into a single tube. 1/4 of the pooled sample was rRNA depleted using Ribo-Zero rRNA removal kit (epicentre). Unbound RNA (rRNA- RNA) was cleaned (using SPRI beads) and reverse transcribed, ligated to linker2 and enriched by PCR as described above (for total PolyA+ libraries). For **digital 3’ gene expression**, library preparation was similar to the total RNA (PolyA+) libraries described above, with one exception- PolyA+ selection was not done before fragmentation, but after linker1 ligation and samples pooling (before the RT reaction step).

#### Western blot analysis

20 fly heads per sample were collected on dry ice. Heads were homogenized in RIPA lysis buffer (50 mM Tris-HCl at pH 7.4, 150 mM NaCl, 1 mM EDTA, 1% NP-40 0.5% Sodium deoxycholate, and 0.1% SDS, 1 mM DTT, protease inhibitor cocktail (Roche) and phosphatase inhibitors) using motorized pestle. After centrifugation lysates were boiled with protein sample buffer (Bio-Rad) and resolved by Criterion XT Bis-Tris gels (Bio-Rad). Antibodies used for western blotting: sheep Anti-mbl antibody was kindly provided by Prof. Darren Monckton (School of Life Sciences, University of Glasgow), mouse anti-tubulin (DM1A; SIGMA, 1:30,000).

#### AGO1-seq procedure

AGO1 immunoprecipitation was performed as described (39, 45). The RNA-seq libraries were performed utilizing the Ovation RNA-seq System for model organisms.

### Physiological and behavioral assessments

#### NMJ isolation from larvae and synaptic die uptake assay

All experiments were carried out at room temperature (22 °C).

Third instar larvae were dissected according to Jan & Jan, 1976, in standard *Drosophila* medium containing 130 mM NaCl, 26 mM sucrose, 5 mM KCl, 2 mM CaCl2, 2 mM MgCl2, 5 mM Hepes, pH 7.4 (46, 47). The preparations were loaded with dye by incubating with 10 μM FM 1-43 in high potassium medium (90 mM KCl and 45 mM NaCl, rest as standard medium) for 30 seconds and subsequently washed for 10 minutes in standard medium without FM dye. Preparations were stimulated by mounting in a platinum plate stimulation chamber (custom made from workshop at Max Planck Institute for Biophysical Chemistry, Güttingen, Germany, 8 mm distance between electrodes) and bathing in standard medium. Stimulation trains of 100 mA were generated using an A310 AccupulserTM triggered by an A385 Stimulus Isolator (both World Precision Instruments, Berlin, Germany). The preparations were stimulated 10 times for 10 seconds at 20 Hz and imaged before unloading and after each stimulation step using an upright Zeiss Axio Examiner.Z1 (Carl Zeiss AG, Oberkochen, Germany) equipped with a 20° 1.0 numerical aperture (NA) water immersion objective (Zeiss) and a 100 W mercury lamp (Zeiss). FM 1-43 fluorescence was detected using a 470/40 HQ excitation filter, a 495 LP Q beam splitter and a 500 LP emission filter (all purchased from AHF, Tübingen, Germany). Images were acquired using a computer operated charge coupled device (CCD) camera (AxioCam MRm, Zeiss). Images were analysed using custom written MATLAB routines (The Mathworks, Inc., Natick, MA, USA).

#### Climbing Assay

We separated both males and females of 3-5 days old three sets of 15 each a day before the experiment is performed.

Three sets of 3-5d old flies separated by sex were placed in 25ml glass graduated cylinders with a mark 10cm from the bottom. We recorded the climbing and plotted the time individual flies need to reach the 10cm mark using 1s as smallest unit.

#### Flight analysis

In both the tapping and free-flight assays we used a Phantom v2012 fast camera (Vision Research, NJ) oriented horizontally and back-illuminated by a near-infrared LED. The camera operated at 10,000 frames per second and resolution of 1280×800 pixels. Triggering was performed manually. In the tapping assay, groups of ~5 files we placed in a bottle with flat sides to allow undistorted imaging. Once the flies climbed to the top of the bottle, we tapped it down against the bench, which caused the flies to fall. We recorded 119 flies during their fall and measured whether they flapped their wings or not. In the free-flight assay we placed an individual fly on a pipette tip or a thin wire and allowed it to take off freely. The fly was released inside a transparent Plexiglas cubic container, with side length of ~20cm, to allow free flight far from any solid boundary. The flapping frequency of 131 flies was manually extracted from the videos.

### Computational analysis

RNA-seq reads were aligned to the genome and transcriptome (dm3) using tophat (48). circRNA detection in RNA-seq data was performed as previously described (12).

CircRNA expression levels were determined by counting back-spliced reads and normalizing to total number of reads. Similarly, for linear RNA expression we used the number of reads from the linear exon-exon junctions from both sides of the circRNA boundaries. samtools-depth tool was used for counting reads within exons.

Differential exon usage analysis was performed using DEXseq (44). The analysis was done using polyA selected library data. UAS-shcircMbl flies were used as a control.

For differential gene expression analysis in circMbl KD with CNS-specific driver (*elav*- *Gal4;UAS-Dcr2)* we used total RNA-seq data. Gene expression levels were determined using HT- seq tool and differential expression analysis was performed with DEseq. Flies expressing shRNA against Luciferase (UAS-shLuc) under the same promoter were used a control. We considered genes with pvalue<0.05 as significantly changing.

Gene expression levels from 3’ DGE experiments were determined using ESAT tool (49) and differential expression analysis was performed with DEseq. We considered genes with fold change>1.5 and pvalue<0.05 as significantly changing. *Actin*-Gal4 flies were used as a control for the lines expressing shRNA under actin promoter. In order to clean non-specific effects, we excluded from downstream analyses genes that were changing in similar direction when comparing the actin-Gal4 control flies and circMbl 8MM KD line. *elav*-Gal4 flies were used as a control for the lines expressing shRNA under *elav* promoter.

topGO package was used for enrichment analysis of gene ontology. GO terms with p-value <0.1 (after FDR correction) were considered significant.

SYLAMER algorithm (38) was used to check for general off-target effect of the shRNA. In order to obtain a list of potential shRNA off target genes we blast all shRNA sequences against the drosophila transcriptome. 3′ RNA-seq data was used to determine the expression level of each putative off-targets gene relative to control line.

For evaluation of circRNAs expression at different developmental stages data was extracted and reanalyzed for circRNA expression as described by (21).

For the analysis of the AGO1-seq we aligned the RNA-seq reads to the genome and transcriptome of *Drosophila* melanogaster (dm6) using STAR (50). Counts per gene were obtained using HTSeq. To assess the distribution of AGO1 immunoprecipitation results we did a Kernel density plot in log2 scale for the mean of normalized gen counts for IP over the normalized gen counts for the input. The AGO1-IP enrichment analysis was done by a negative binomial GLM approach using DeSeq2 (51). For this analysis we compared INP an IP counts for control and sh-circMbl flies indepently. To further compare these results from both control and sh-Mbl we did a Likelihood Ratio Test (LTR) comparing a simpler model that only considers IP and input as factors with another more complex that includes also an interaction between the IP results and the genetic background.

To see possible sh-circMbl effects, we analyzed 6nt kmer enrichment in the 3′UTR of the genes using Sylamer algorithm (38). For this, we ranked the gene list by Log2 fold change multiplied by the inverse of the p adjusted value (log2FC*1/(pval)) the results of the IP enrichment analysis.

