## Supplementary material for "An *in vivo* knockdown strategy reveals multiple functions for circMbl"

**Figure Legends to Figures S1-S4**

**Figure S1. Generating and validating a resource for circRNA knockdown. A.** shRNAs can be used to downregulate a circRNA expressed from a transfected plasmid. *Drosophila* S2 cells were transfected with a circMbl expressing plasmid (pMT-circMbl) with or without a plasmid expressing a shRNA against the circMbl backsplicing junction. (n=3; Student`s t-test; **p=0.01). **B.** AGO-IP enriched genes contains conserved miRNA binding sites. The x-axis represents the genes sorted from the most to the least enriched in the AGO-1 IP-seq. The y-axis shows the percentage of genes with more than 1 conserved miRNA binding site for each group of 30 genes. Left panel WT and right sh-line is the non-parametric local regression of the data with the confidence area. **C.** Sylamer enrichment landscape plot for sh-circMbl and sh-circMbl* 6mers. The x-axis represents the genes sorted by adjusted pvalue in the interaction analysis. The y-axis shows the hypergeometric significance for each word at each leading bin.

**Figure S2. Modulation of circMbl in flies.** Gene Ontology (GO) terms enriched in circMBL KD vs. circMBL over expression lines using the actin-Gal4 driver. For each line, list of differentially expressed genes obtained from the 3' RNA-seq analysis (relative to actin-Gal4 control) was used for GO terms enrichment analysis. In order to clean non-specific effects, we excluded from the analysis genes that were changing in similar direction when comparing the actin-Gal4 control flies and circMbl 8MM KD. Top GO package in R was used to determine the statistical significance of the enriched group. The data presented is p-value after FDR correction.

**Figure S3. Knockdown of circMbl using different drivers. A.** Downregulation circMbl KD using CNS-specific driver elav-Gal4 with and without co-expression of Dicer-2. Expression levels were determined from total RNA-seq data. The number of back-spliced reads (for circRNA expression) or linear exon-exon junction reads from both sides of the circRNA boundaries (for mRNA expression) were counted and normalized to the total number of reads (n=3).

**Figure S4. sh-RNA shift mutants partially recapitulate circMbl-KD phenotype. A.** Expression levels of circMbl and mbl mRNA in flies expressing the indicated shRNAs against circMbl. Data was normalized to rp49 ($n \geq 4$; Student`s t-test; $^{**}p<0.01$, $^{***}p<0.0005$). **B.** males and **C.** females of circMbl KD, showed delayed climbing when compared to actin-Gal4 controls. P values were obtained by comparing each strain with flies carrying the actin-Gal4 (n=15x9 sets, Student`s t-test; $^{*}p<0.05$, $^{**}p<0.01$, $^{***}p<0.0005$). **D.** Knockdown of circMbl using the UAS-shcircMbl KD3 transgene and the actin-Gal4 driver show distinctive climbing defect when compared to control flies (circMbl 8MM flies: actin-Gal4, UAS-shcircMbl 8MM). Graph shows the average of 5 vials of flies (20 flies/vial; Student`s t-test; $^{*}p<0.05$, $^{**}p<0.01$). **E.** Histogram of the mean wing-beat frequency of circMbl lines measured in the free-flight assay ($n_{males}$=25/33, $n_{females}$=29/12/32; Student`s t-test; $^{*}p<0.05$, $^{***}p<0.0005$).

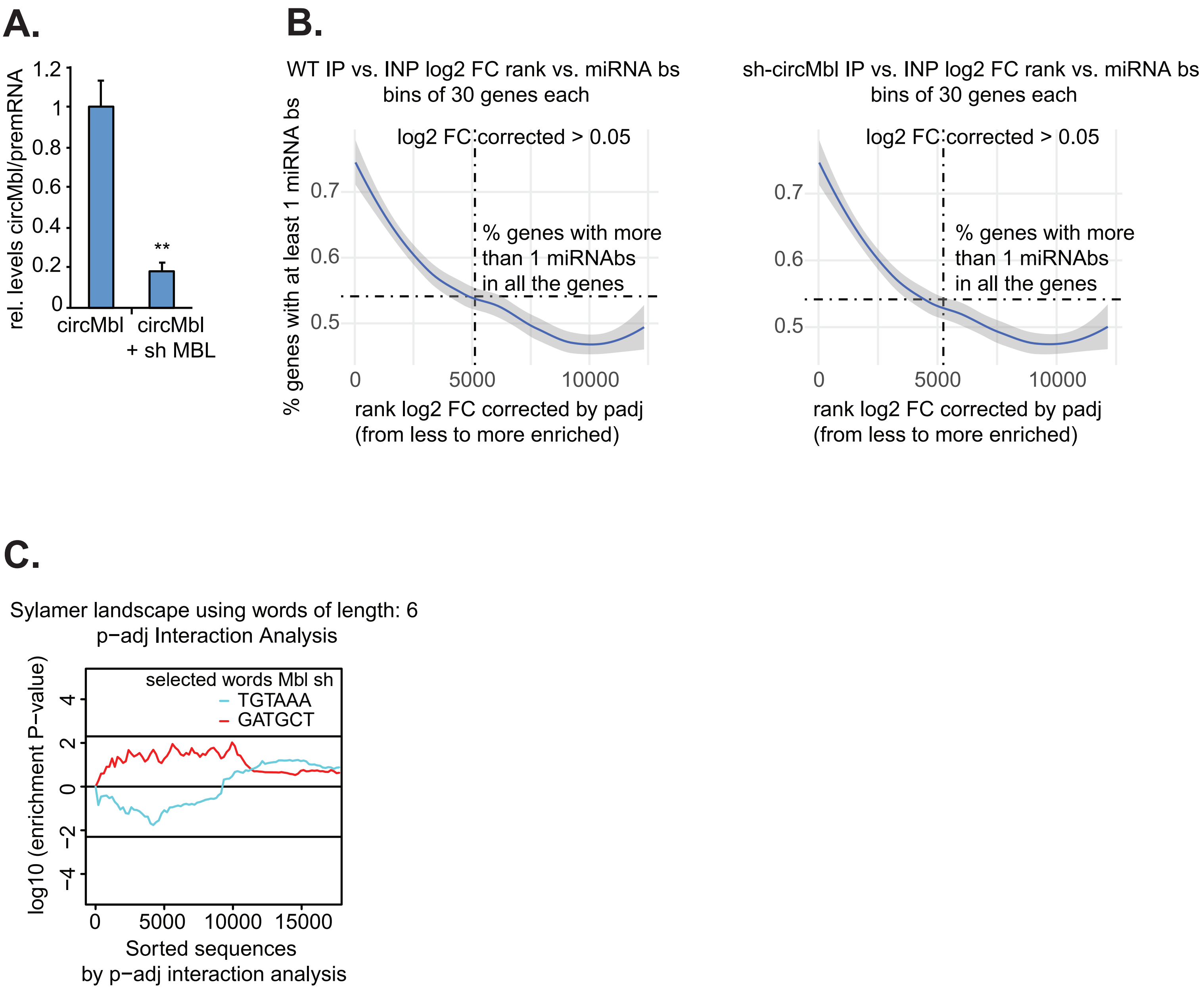

A.
rel. levels circMbl/premRNA
1.2
1
0.8
0.6
0.4
0.2
0
**
circMbl
circMbl + sh MBL
B.
WT IP vs. INP log2 FC rank vs. miRNA bs
bins of 30 genes each
log2 FC corrected > 0.05
% genes with more than 1 miRNAbs in all the genes
% genes with at least 1 miRNA bs
0.7
0.6
0.5
0
5000
10000
rank log2 FC corrected by padj (from less to more enriched)
sh-circMbl IP vs. INP log2 FC rank vs. miRNA bs
bins of 30 genes each
log2 FC corrected > 0.05
% genes with more than 1 miRNAbs in all the genes
0.7
0.6
0.5
0
5000
10000
rank log2 FC corrected by padj (from less to more enriched)
C.
Sylamer landscape using words of length: 6
p−adj Interaction Analysis
selected words Mbl sh
TGTAAA
GATGCT
log10 (enrichment P−value)
4
2
0
−2
−4
0
5000
10000
15000
Sorted sequences by p−adj interaction analysis


**Figure S1**

| GO.ID | Term | circMbl KD | circMbl OE |
|---|---|---|---|
| GO:0008010 | structural constituent of chitin-based l... | 1.23256E-07 | 2.5823E-12 |
| GO:0005576 | extracellular region | 6.1628E-07 | 0.004557 |
| GO:0031012 | extracellular matrix | 0.00028613 | 6.8355E-13 |
| GO:0005859 | muscle myosin complex | 0.00028613 | 0.003764478 |
| GO:0040003 | chitin-based cuticle development | 0.000360964 | 1.80002E-12 |
| GO:0005615 | extracellular space | 0.026412 | 0.079077353 |
| GO:0015986 | ATP synthesis coupled proton transport | NA | 2.62028E-10 |
| GO:0046933 | proton-transporting ATP synthase activit... | NA | 1.45824E-08 |
| GO:0005811 | lipid particle | NA | 1.6709E-08 |
| GO:0005747 | mitochondrial respiratory chain complex ... | NA | 8.463E-08 |
| GO:0006030 | chitin metabolic process | NA | 5.58233E-07 |
| GO:0006120 | mitochondrial electron transport, NADH t... | NA | 4.05067E-06 |
| GO:0008540 | proteasome regulatory particle, base sub... | NA | 5.0127E-06 |
| GO:0008061 | chitin binding | NA | 5.38555E-06 |
| GO:0000276 | mitochondrial proton-transporting ATP sy... | NA | 1.29115E-05 |
| GO:0000275 | mitochondrial proton-transporting ATP sy... | NA | 1.75269E-05 |
| GO:0008121 | ubiquinol-cytochrome-c reductase activit... | NA | 0.00014973 |
| GO:0008137 | NADH dehydrogenase (ubiquinone) activity | NA | 0.0001519 |
| GO:0055114 | oxidation-reduction process | NA | 0.000236394 |
| GO:0006122 | mitochondrial electron transport, ubiqui... | NA | 0.000294865 |
| GO:0005750 | mitochondrial respiratory chain complex ... | NA | 0.000329117 |
| GO:0004129 | cytochrome-c oxidase activity | NA | 0.001295147 |
| GO:0043161 | proteasome-mediated ubiquitin-dependent ... | NA | 0.00164052 |
| GO:0005751 | mitochondrial respiratory chain complex ... | NA | 0.0021049 |
| GO:0016490 | structural constituent of peritrophic me... | NA | 0.003314182 |
| GO:0006123 | mitochondrial electron transport, cytoch... | NA | 0.00947856 |
| GO:0007629 | flight behavior | NA | 0.017176385 |
| GO:0009055 | electron carrier activity | NA | 0.021941111 |
| GO:0042559 | pteridine-containing compound biosynthet... | NA | 0.03255 |
| GO:0030018 | Z disc | NA | 0.034570345 |
| GO:0004298 | threonine-type endopeptidase activity | NA | 0.04557 |

**Figure S2**

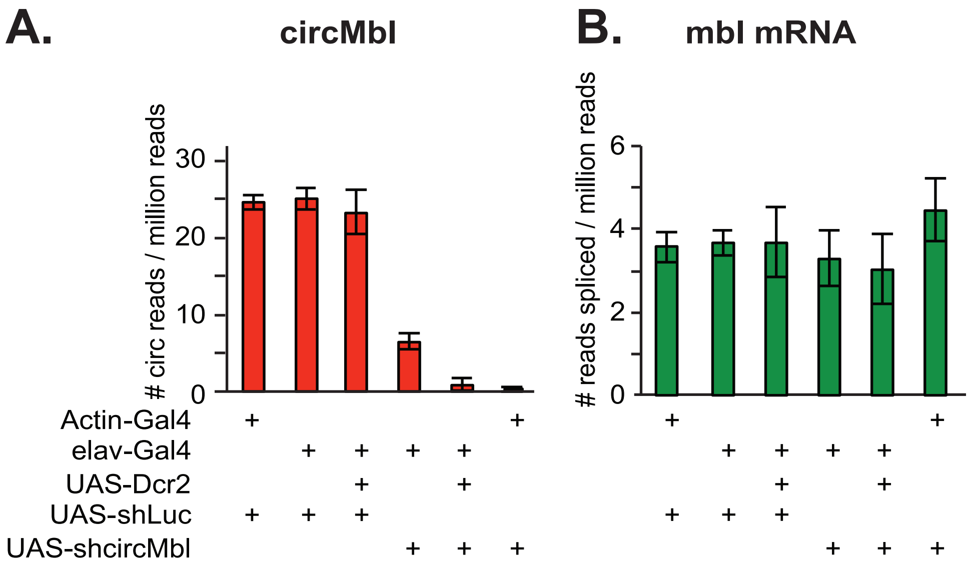
A.
circMbl
### circ reads / million reads
30
20
10
0
Actin-Gal4
elav-Gal4
UAS-Dcr2
UAS-shLuc
UAS-shcircMbl
B.
mbl mRNA
### reads spliced / million reads
6
4
2
0

### Figure S3

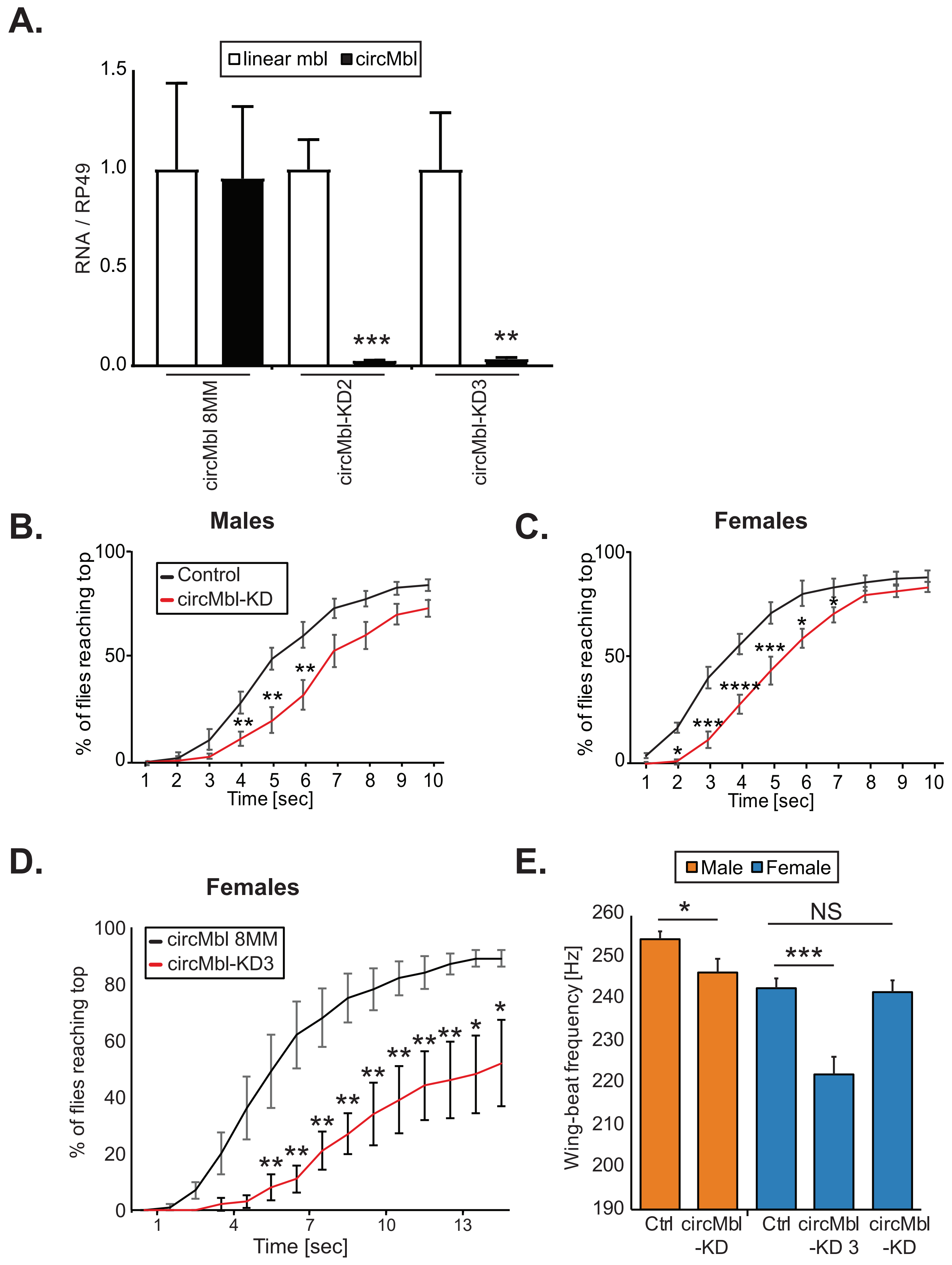
A.
linear mbl
circMbl
RNA / RP49
1.5
1.0
0.5
0.0
***
**
circMbl 8MM
circMbl-KD2
circMbl-KD3
B.
Males
Control
circMbl-KD
% of flies reaching top
Time [sec]
C.
Females
% of flies reaching top
Time [sec]
D.
Females
circMbl 8MM
circMbl-KD3
% of flies reaching top
Time [sec]
E.
Male
Female
Wing-beat frequency [Hz]
*
NS
***
Ctrl
circMbl -KD
Ctrl
circMbl -KD 3
circMbl -KD

Figure S4
